# Discovery of *Plasmodium falciparum* SR12 as a GOLD-Domain seven transmembrane protein regulating GPCR trafficking in mammalian cells

**DOI:** 10.1101/2020.04.17.047217

**Authors:** Pedro H. S. Pereira, Sajjad Ahrari, Camila L. Kiyan, Hiroyuki Kobayashi, Miriam S. Moraes, Michel Bouvier, Celia R. S. Garcia

## Abstract

Considered a significant public health issue, the growing resistance to conventional antimalarials necessitates the identification of new targets for drug development. Given that G protein-coupled receptors (GPCRs) are readily druggable targets, we explored the cellular role and potential structure of a GPCR-like protein identified in the *P. falciparum* genome, serpentine receptor 12 (SR12). Alphafold structure analysis, coupled with molecular dynamics simulations of SR12, revealed structural similarities to the Golgi dynamics domain (GOLD)-seven-transmembrane helix protein family (GOST proteins). This family of proteins, which includes TMEM87A and the orphan GPCRs GPR180, GPR107, and GPR108, is involved in subcellular trafficking. Consistent with such a trafficking role, SR12 is mainly present in the secretory pathway when expressed in mammalian cells. Co-expression of SR12 with GPCRs PAR1 and M3R led to increased plasma membrane targeting of these receptors. SR12 expression in HEK293 cells conferred Gαq-dependent calcium signaling in response to the protease activated receptor 1 (PAR1) agonist thrombin. This response was completely abrogated in cells genetically devoid of PARs (PAR KO cells), consistent with its functions as a chaperone-like protein, promoting receptor trafficking to the plasma membrane. Taken together, the data show that the *Plasmodium falciparum* SR12 promotes GPCR trafficking when expressed in mammalian cells. Although the physiological consequences of such activity remain to be determined, the finding revealed the presence of a GOST protein in the parasite genome.

## Introduction

Malaria, a disease caused by the *Plasmodium* genus protozoa, constitutes significant global health concern, with *P. falciparum* being responsible for the most severe cases. Approximately half of the world population inhabits regions at risk of infection, resulting in 263 million reported cases and over 597,000 deaths worldwide (1). The widespread resistance of parasites to drugs such as chloroquine and pyrimethamine/sulfadoxine is already a global issue. Additionally, reports of resistance to the latest drugs, such as artemisinin, atovaquone, and piperaquine, are on the rise, particularly in Southeast Asia. Consequently, the need for alternative strategies to eradicate the disease is becoming increasingly urgent (2, 3).

Remarkably, parasites have demonstrated the ability to sense external molecules. *Plasmodium*, for instance, has been reported to possess at least four G protein-coupled receptors (GPCRs)-like proteins based on predicted membrane topologies: SR1, SR10, SR12, and SR25 (4). Among these receptors, SR25 has been identified as a potassium sensor, linked to Ca^2+^ signaling via a PLC-dependent pathway. This receptor modulates stress survival and antimalarial action in parasites (5).

SR12 shares 30% primary sequence homology with the human intimal thickness-related receptor GPR180 associated with vascular restenosis (6). GPR180 may recognize lactate (7) and has been proposed to be part of the TGFβ signaling complex (8). It has also been shown to be essential for adeno-associated virus transduction (9, 10). Recently, it was found to localize primarily intracellular vesicles and to participate in cargo transport (11). In addition to GPR180, SR12 shares structural similarities with human orphan GPCRs and seven-transmembrane proteins, including GPR107, GPR108, and TMEM87A. GPR107 and GPR108 share 28% similarity with SR12, are localized in the Golgi, and implicated in subcellular transport (9, 10, 12, 13). The three-dimensional structure of TMEM87A has recently been elucidated, describing a Golgi Dynamics domain (GOLD) in its N-terminal region fused to a seven-transmembrane (7TM) bundle, supporting its role in the subcellular trafficking of membrane-associated cargoes, and classified as a GOLD-seven-transmembrane helix protein (GOST proteins) family member (14). The orphan GPCRs GPR107, GPR108 and GPR180 have also been suggested to be part of the GOST protein family. Despite having only one transmembrane domain, and thus not being classified as a GOST, p24A is a GOLD containing protein that was shown to interact and control PAR2 trafficking (15).

GPCRs form the largest transmembrane protein family involved in signal transduction across biological membranes (16, 17). In humans, approximately 800 members were identified, half of which are involved in sensory functions such as smell, taste, and light perception (18). GPCRs are also found throughout evolution in species as diverse as yeast, flagellated protozoa and unicellular protists. Structurally, GPCRs share a common architecture with seven hydrophobic transmembrane domains connected by extracellular and intracellular loops, an extracellular N-terminus, and an intracellular C-terminus. Signaling through GPCRs is initiated with agonist binding from the extracellular side, subsequently transmitting signals to heterotrimeric G proteins (19, 20). Despite the 7TM topology that resembles a GPCR, SR12 lacks all other canonical motifs described in GPCRs able to signal through G-proteins, such as DRY, PIF and NPY motifs. This, taken together with the fact that no heterotrimeric G proteins have been identified to date in *Plasmodium,* suggests that the seven-transmembrane-domain protein found in the parasite may have functions different from those of canonical GPCRs (21, 22). Interestingly, some seven transmembrane-containing proteins were found not to have G protein signaling functions (23, 24).

Given the GPCR-like membrane topology of SR12 and its similarity with other orphan GPCRs/GOLD-domain containing proteins involved in protein trafficking (11, 14), we set out to assess the potential role of SR12 in either signaling or cargo transport as well as its structural similarity with GOST proteins. Our findings align with the inferred function of the GOLD-domain seven-transmembrane helix protein family, thereby classifying SR12 as a member of this group and supporting a role in GPCR trafficking.

## Results

### Molecular modeling of SR12 structure

Based on the principle that structure underlies function, we modeled the putative SR12 structure using AlphaFold2 (AF2) and AlphaFold3 (AF3). Both models adopt a similar overall architecture (Figure 1A, B) and exhibit a predicted local distance difference test score (pLDDT) > 70 across most residues 100–460 (Figure 1C), indicating high confidence in the predicted local geometry. Although both versions of AF have high confidence scores, AF3 assigns higher pLDDT values than AF2 in the 100–120 region (Figure 1C). In both models, SR12 comprises an N-terminal helical region (residues 1–88) with very low pLDDT, yielding uncertainty on that part of the structure. However, greater pLDDT for the two models predict, with high confidence, a globular domain enriched in β-strands (residues 88–230), and a C-terminal helical domain (residues 231–470) containing seven transmembrane helices and a short helical appendage reminiscent of GPCR helix 8, yielding an overall GPCR-like fold (Figure 1D). Least-squares superposition of AF2 and AF3 indicates that the C-terminal transmembrane region is the most structurally conserved between models (Figure 1E and S1). The globular domains also share a similar fold but are positioned closer to the 7TM bundle in AF2 than in AF3. In contrast, the N-terminal helical region adopts divergent orientations in the two models (Figure S1), consistent with its low pLDDT and correspondingly low confidence in its placement.

**Figure 1.**
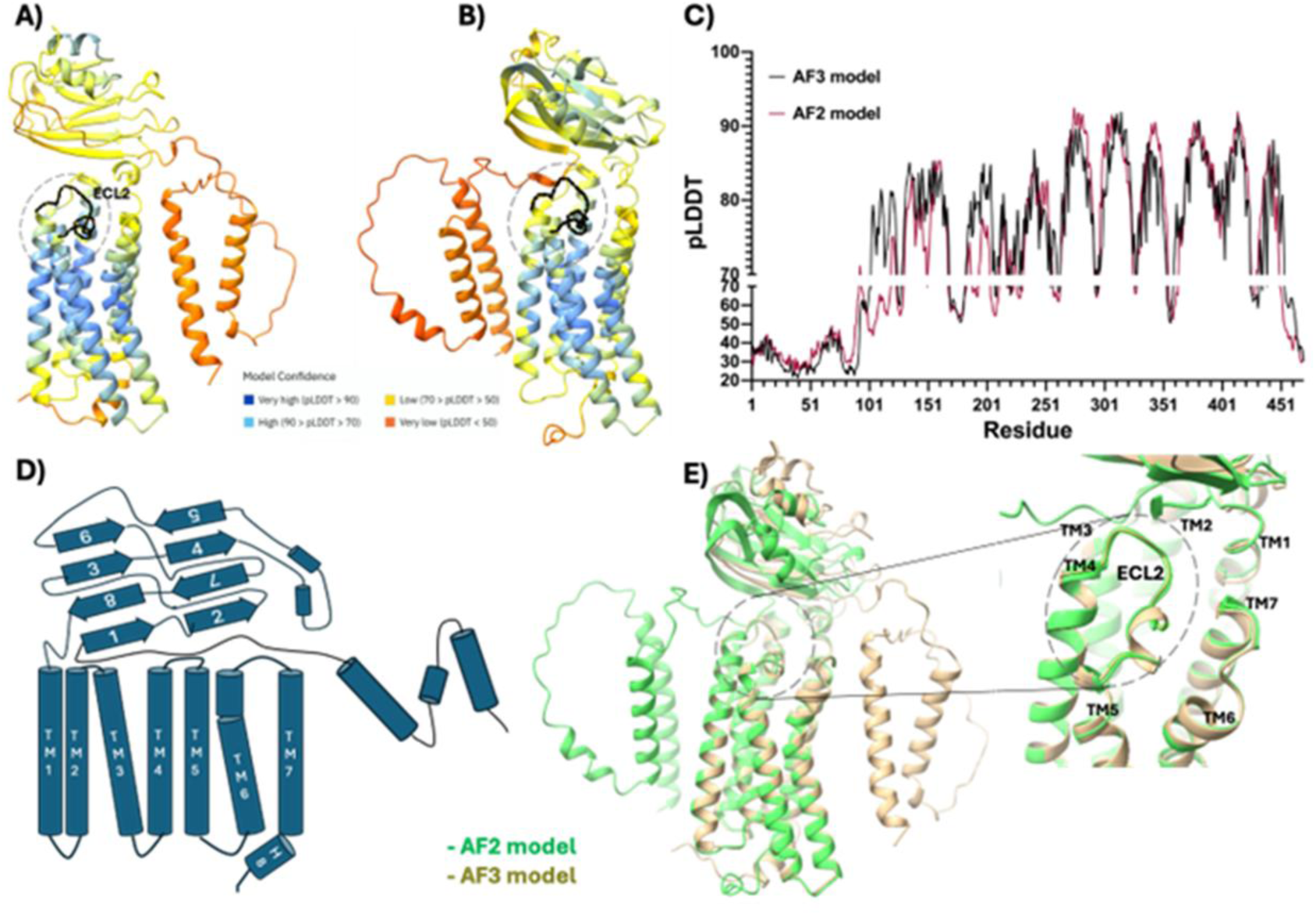
AlphaFold modeling of SR12. (**A**, **B**) Three-dimensional structures predicted by AF2 and AF3, colored by per-residue pLDDT. (**C**) Per-residue pLDDT profiles for AF2 (red) and AF3 (black). (**D**) Schematic of the SR12 amino-acid sequence and domain organization. (**E**) Least-squares superposition of AF2 (green) and AF3 (tan) models.

We further assessed the integrity and conformational stability of the two models by molecular dynamics (MD) simulations in a native-like environment. Although the overall 3D fold was retained, the β-strand content of the globular domain changed in both models (Figure 2). Each model initially contained six β-strands; during simulation, low-pLDDT regions tended to lose β-strand character and adopt coils or bends, whereas high-pLDDT regions largely preserved their predicted secondary structure (Figure 2A, B). A similar trend was observed for the 7TM region (Figure 2C, D). Notably, AF3 pLDDT values more accurately reflected secondary-structure stability during simulation. In the AF2 trajectory, the globular domain evolved to eight β-strands and two α-helices. In contrast, the AF3 trajectory retained seven stable β-strands and two α-helices, with the first β-strand disappearing midway through the simulation.

**Figure 2.**
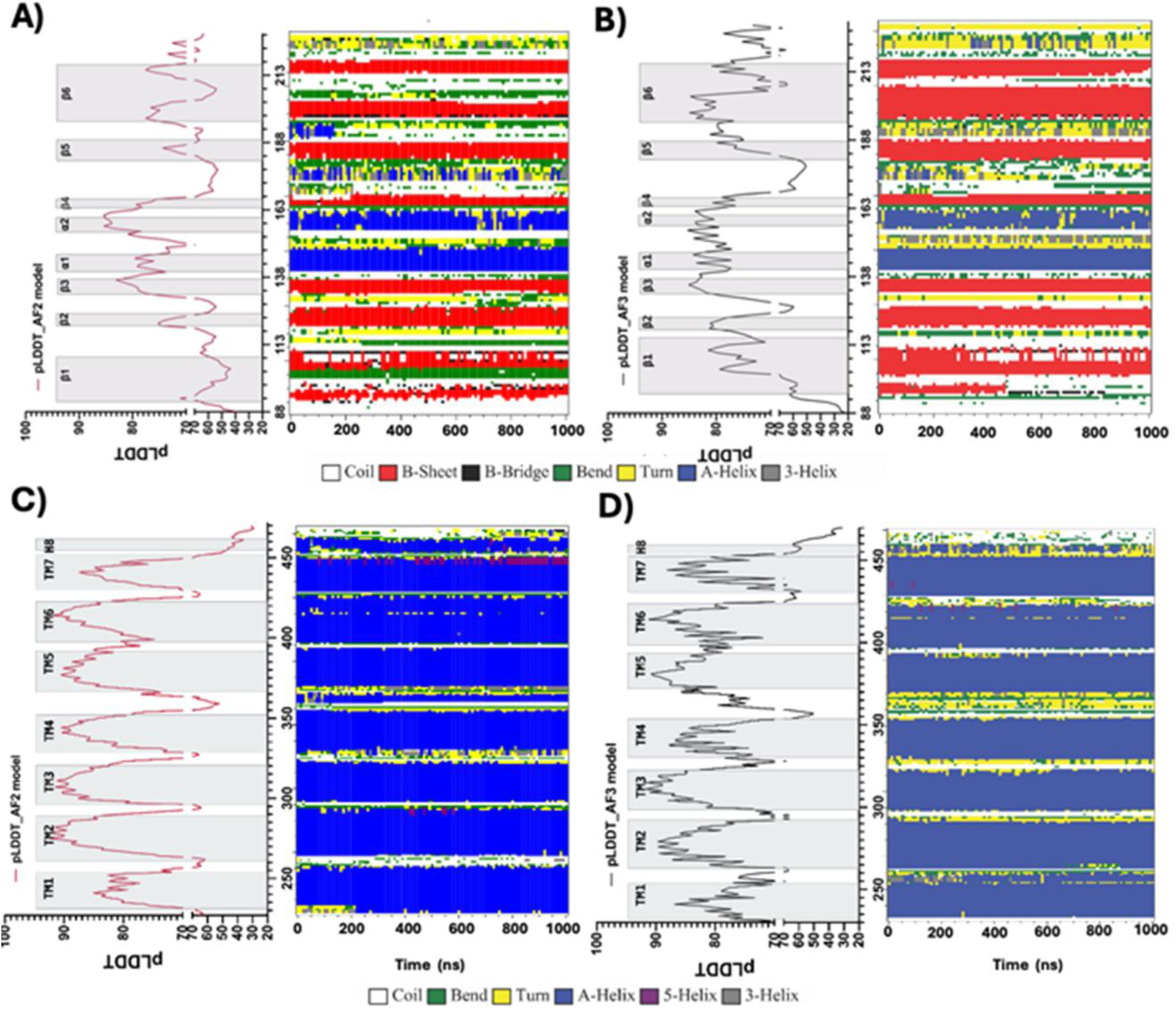
Stability of AlphaFold-predicted secondary structure in SR12 during 1 µs MD simulations. The globular domain (top) and 7TM region (bottom) are shown. (**A**, **B**) Per-residue pLDDT profiles and AlphaFold-predicted secondary structure are presented together with the MD-derived secondary-structure timeline (DSSP) for the globular domain. (**C**, **D**) The corresponding pLDDT, AlphaFold secondary-structure prediction, and DSSP timeline are shown for the 7TM region. For each panel, AF2 is shown on the left and AF3 on the right.

### GPR180 is the closest structural homolog of SR12

The AF3-derived SR12 model was queried against the Foldseek server to identify structural homologs. Twenty-seven high-confidence hits (E-value < 10⁻⁶) were retained and clustered into three groups based on pLog(E-value) (Figure 3A). The top matches were predominantly orthologs of GPR180, TMEM145, TMEM87A, and TMEM87B. These proteins share a common architecture consisting of a large N-terminal Golgi dynamics (GOLD) domain fused to a seven-transmembrane (7TM) bundle and are classified within the GOST proteins family. LUSTR/GOST proteins have been reported to localize to the secretory pathway, including the ER and Golgi (9, 10, 12, 25–30).

**Figure 3.**
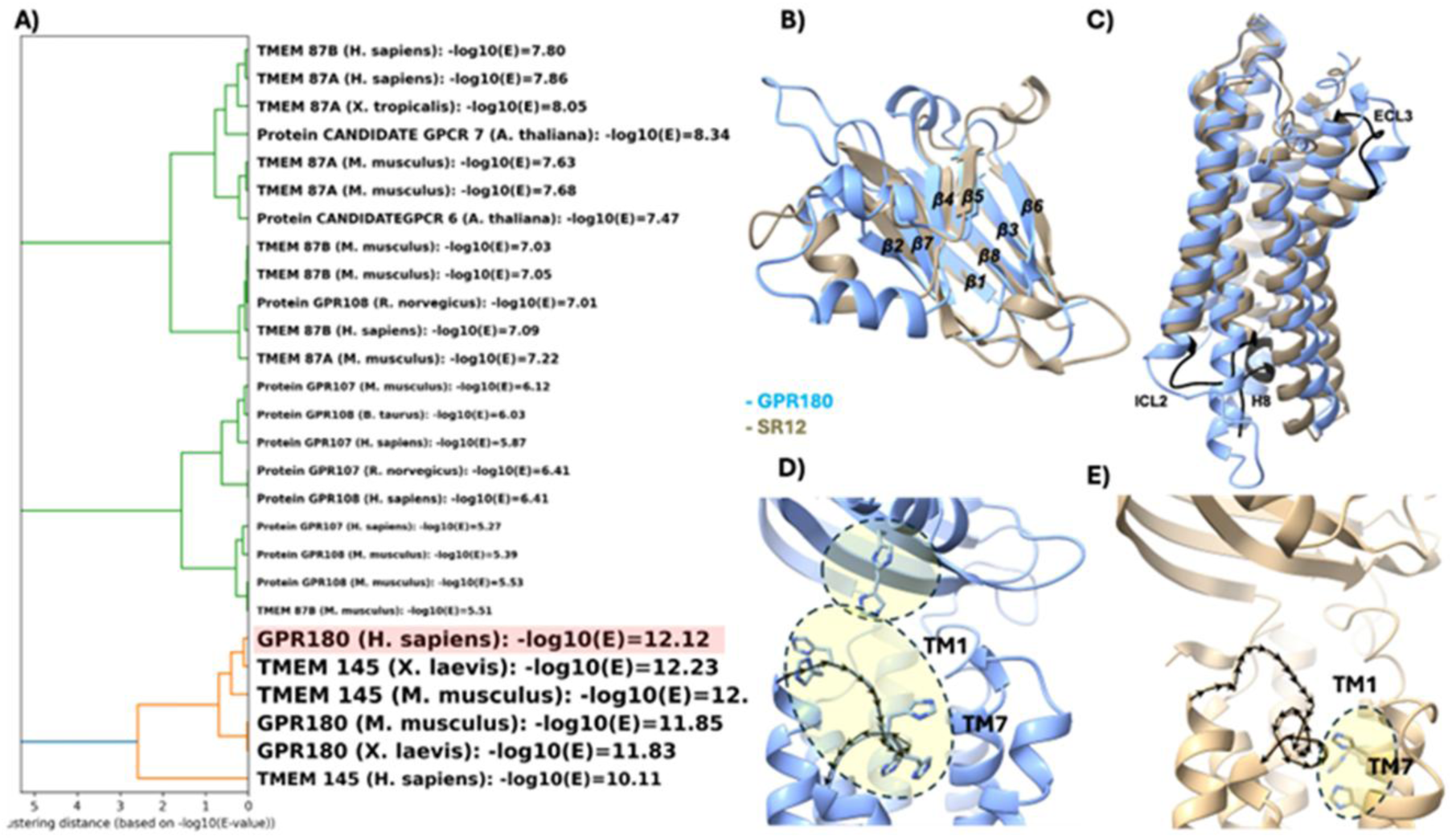
Structural comparison of SR12 and GPR180. (**A**) Clustering of the closest SR12 structural homologs identified by Foldseek based on E-values. (**B**, **C**) Superposition of the N-terminal domain and 7TM bundle of GPR180 (blue) with SR12 (tan) both generated by AF3. The GPR180 N-terminal GOLD domain for its part was taken from the resolved mouse crystal structure (PDB: 9FOW) of this domain from GPR180. (**D**) Histidine-rich patch in GPR180, located on ECL2 and the GOLD domain. (**E**) Corresponding histidine positions in SR12 are mapped onto the 7TM domain.

Among the highest-scoring homologs was human GPR180 (pLog(E-value) = 12.1; Figure 3A). The recently resolved crystal structure of the mouse GPR180 N-terminal globular domain (11) exhibited strong structural similarity to the SR12 N-terminal globular region predicted by AF3 (Figure 3B). This domain corresponds to a GOLD domain and adopts a β-sandwich fold comprising seven to eight β-strands arranged into two opposing sheets, with several short α-helices decorating strand-connecting loops. This β-sandwich topology is conserved among GOLD domains, whereas the lengths and conformations of flanking loops vary across homologs (11). Superposition of the GPR180 GOLD domain crystal structure (PDB: 9FOW) onto SR12 shows close agreement in β-strand number and placement (Figure 3B), supporting assignment of the SR12 N-terminal globular region as a GOLD domain. The main differences were the apparent absence of the α1/α2 helices in the SR12 model and variations in loop lengths.

The SR12 7TM region also aligned closely with the GPR180 7TM bundle, indicating high similarity among the transmembrane helices organization (Figure 3C). Differences were primarily confined to intracellular loop 2 (ICL2) and helix 8 (H8). In addition, the location of the histidine-rich patch differed between the two proteins (Figures 3D and 3E). In GPR180, the histidine cluster on ECL2 and within the GOLD domain has been implicated in pH-dependent conformational changes that alter the relative orientation of the GOLD and 7TM domains (11) contributing to its cargo trafficking function. In the SR12 model, the corresponding histidines were instead positioned on the extracellular faces of TM1 (H237) and TM7 (H432, H434, H438) (Figure 3E).

### SR12 is located mainly in subcellular organelles

Both GPR180 and TMEM67A, as well as the orphan GPCRs GPR107 and GPR108, are proposed to be Golgi-resident and to participate in endosomal protein transport. To examine whether the subcellular localization pattern of SR12 aligns with its proposed function, we employed ebBRET assays using C-terminally rLuc-tagged SR12, and rGFP constructs directed to distinct subcellular compartments through fusion with organelle-specific targeting sequences. These compartments included the plasma membrane (PM), early endosomes (EE), Golgi apparatus (Golgi), mitochondria (Mito), and endoplasmic reticulum (ER). Although mitochondria is not canonically related to trafficking to the plasma membrane, GOLD-containing proteins are found in this organelle, such as ACBD3, related to cholesterol transport (31). When compared with *bona fide* GPCRs such as the protease activated receptor (PAR) 1 and muscarinic receptor 3 (M3R), although SR12 could reach the plasma membrane, it was proportionally found in more abundance in the endoplasmic reticulum and mitochondria (Figure 4A).

**Figure 4.**
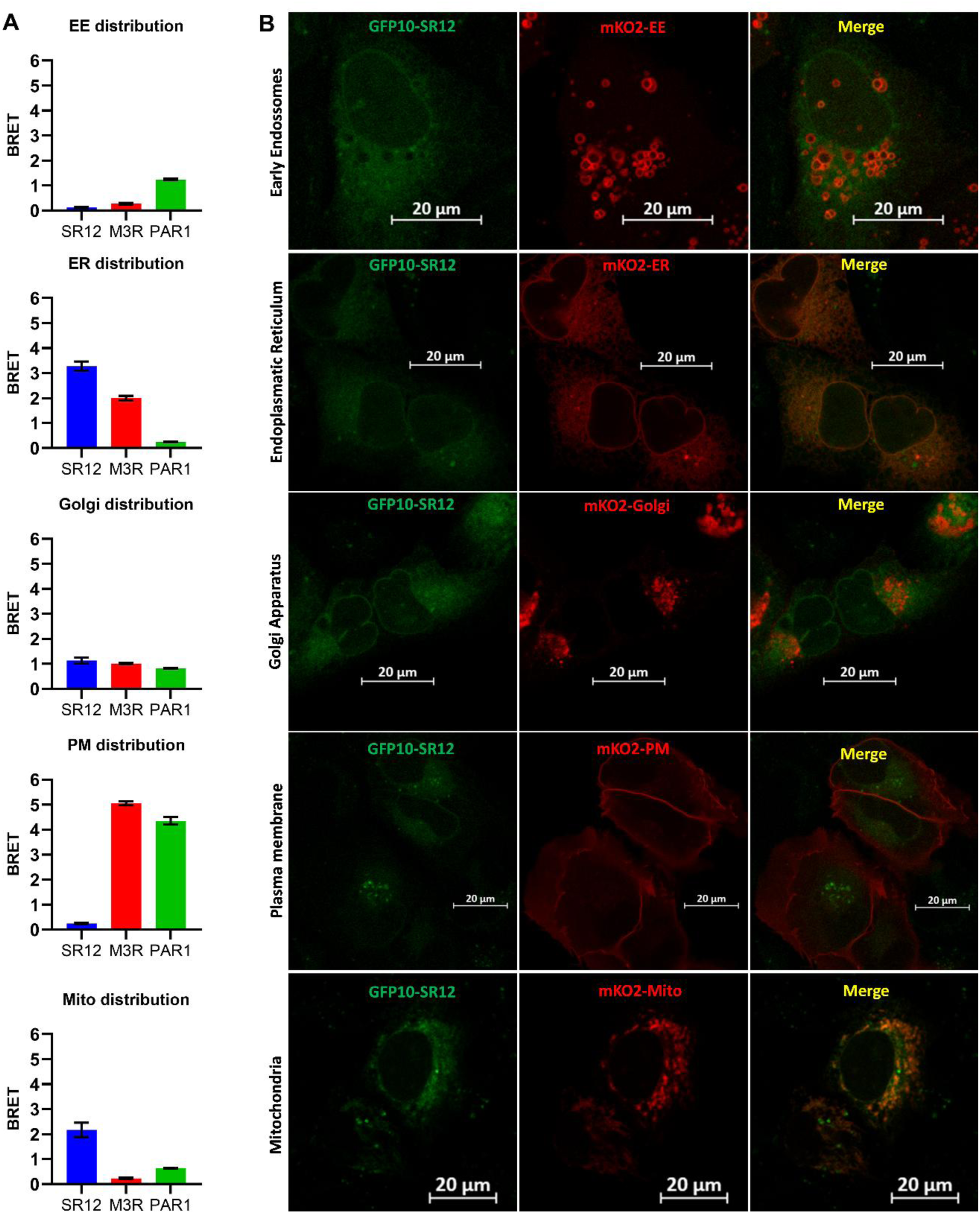
Subcellular localization of SR12. (**A**) HEK293 Cells were co-transfected with C-terminally rLuc tagged full-length SR12 and with rGFP constructs directed to subcellular compartments such as plasma membrane (PM), early endosomes (EE), Golgi apparatus (Golgi), endoplasmic reticulum (ER), and mitochondria (Mito). BRET values were calculated by dividing the intensity of light emitted by rGFP (515 nm) by the intensity of light emitted by rLuc2 (400 nm). Values were acquired without stimulation, and results are expressed as raw BRET values (mean ± SEM, n=3). (**B**) U2OS cells were co-transfected with C-terminally GFP10-tagged full-length SR12 and with mKO constructs directed to subcellular compartments such as plasma membrane (PM), early endosomes (EE), Golgi apparatus (Golgi), endoplasmic reticulum (ER), and mitochondria (Mito), and images were acquired by confocal microscopy.

Confocal microscopy experiments using a GFP10-tagged SR12 and mKO-tagged proteins targeted to specific organelles largely confirmed the BRET data (Figure 4B). The images show colocalization of SR12-GFP10 with both Golgi and ER markers. In contrast with the BRET data that revealed weak localization at the plasma membrane and early endosomes, the microscopy data could not detect a convincing expression of SR12 in these organelles. Of note, we also detected the presence of SR12 in small round structures inside the cells that do not match any of the fluorescent markers used in this experiment. The nature of this structure remains to be identified.

### Protonation of lumen exposed histidines perturbs SR12 conformation and expands the extracellular pocket

Because histidine side chains can become protonated under acidic conditions, they are expected to be partially protonated in acidified intracellular compartments where GOST proteins have been found to reside. To test the structural impact of this state on SR12, the luminal exposed histidines (that can face either the extracellular or the lumen of organelles compartments) were protonated to mimic a plausible positively charged form. Backbone RMSD analyses during MD indicated that histidine protonation induced larger deviations from the starting structure than the non-protonated condition (Figure 5A). Consistently, RMSF profiles showed increased flexibility in the GOLD domain and in ECL1 (TM2–TM3) and ECL2 (TM4–TM5) upon histidine protonation (Figure 5B).

**Figure 5.**
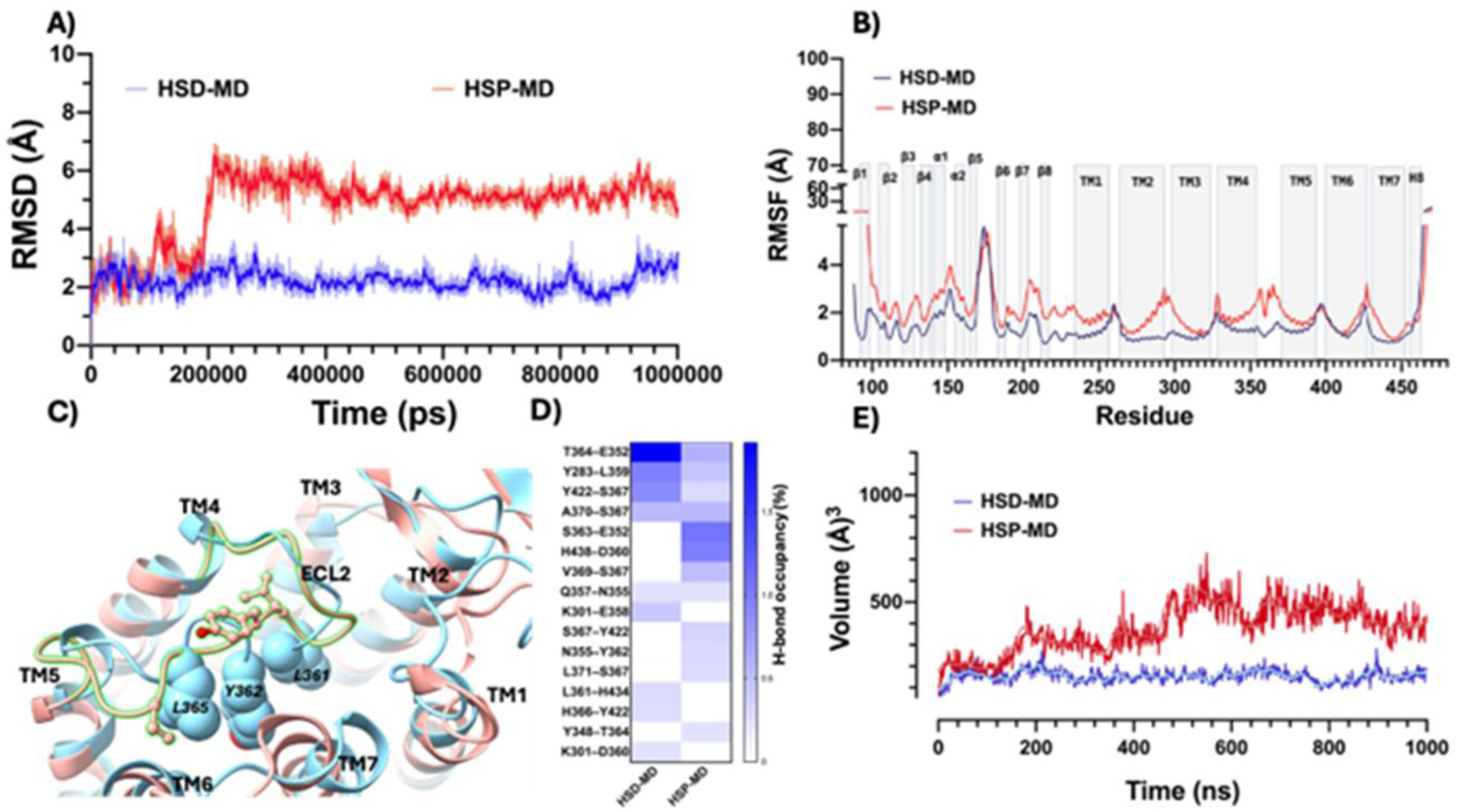
Protonation of extracellularly exposed histidines increases conformational drift and expands the SR12 extracellular pocket. (**A**) Time evolution of backbone RMSD for simulations with neutral histidines (HSD; blue) and protonated histidines (HSP; red). (**B**) Per-residue RMSF for HSD (blue) and HSP (red), highlighting increased mobility in the GOLD domain and extracellular loops. (**C**) Representative snapshots from HSD (blue) and HSP (red) trajectories illustrating ECL2 reorientation. (D) Protonation-dependent remodeling of the ECL2 hydrogen-bond network, summarized as changes in hydrogen-bond occupancy across the trajectories. (**E**) Extracellular pocket volume as a function of time, showing pocket expansion in HSP (red) relative to HSD (blue).

These perturbations were accompanied by a pronounced rearrangement of ECL2, particularly involving Leu361, Tyr362, and Leu365 (Figure 5C). In the AF3 model, these bulky residues projected into the opening of the 7TM bundle facing the GOLD domain and partially occluding it. Upon histidine protonation, they rotated away, thereby exposing the pocket (Figure 5C). Concordantly, the hydrogen-bond network connecting ECL2 to the rest of the protein was extensively remodeled (Figure 5D). Together, these changes increased the volume and accessibility of 7TM bundle opening, potentially favoring ligand or cargo binding (Figure 5E). A second, independent set of MD simulations reproduced these effects, confirming protonation-dependent structural perturbations, ECL2 reorientation, and pocket expansion (Figure S2).

### SR12 increases the localization of PAR1 and M3R in the secretory system

To investigate the impact of SR12 on the expression and/or localization of canonical GPCRs, we assessed whether the co-expression of SR12 with M3R-rLuc or PAR1-rLuc could induce alterations in subcellular localization. BRET between PAR1-rLuc or M3R-rLuc and rGFPs targeted to specific subcellular compartment membranes revealed that the co-expression of SR12 led to PAR1 and M3R increase in abundance, respectively, of 25% and 39% in recycling endosomes (RE), 18% and 28% in early endosomes, 18% and 20% in the Golgi apparatus, as well as 13% and 15% on the plasma membrane (Figure 6). The BRET signal between the receptors and the ER marker was not affected by SR12 expression. *Plasma membrane* BRET results were further validated using ELISA and HA-tagged receptors, in which co-expression of SR12 and M3R or D2R increased the surface expression of these GPCRs (Supplementary Figure S3). SR12 can therefore affect the intracellular distribution of receptors, increasing the abundance of PAR1 and M3R on the Golgi apparatus, plasma membrane, and endosomal vesicles.

**Figure 6.**
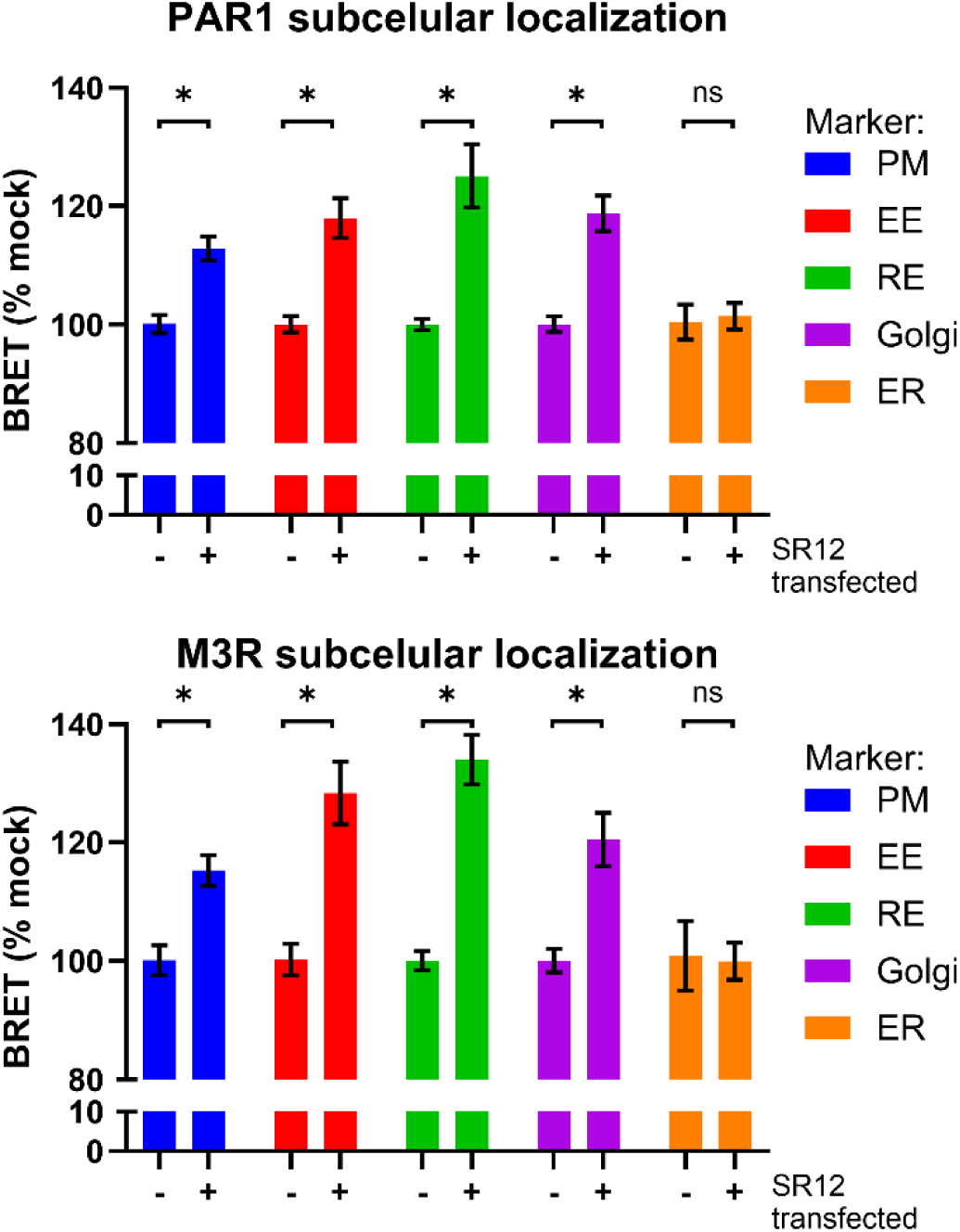
Subcellular localization of PAR1 and M3R in presence of SR12. HEK293 Cells were co-transfected with C-terminally rLuc tagged full-length PAR1 (left) or M3R (right), untagged SR12, and with rGFP constructs directed to subcellular compartments such as plasma membrane (PM), early endosomes (EE), Golgi apparatus (Golgi), endoplasmic reticulum (ER), and recycling endosomes (RE). BRET values were calculated by dividing the intensity of light emitted by rGFP (515 nm) by the intensity of light emitted by rLuc2 (400 nm), and results were normalized and expressed as percentage of mock transfected cells (mean ± SEM, n=3). * Statistically significant by two-way ANOVA with Šidák post-test, p<0,05

### SR12 promotes signaling of endogenously expressed receptors

Given the effect of SR12 expression on PAR1 subcellular localization and the role of single TM GOLD-containing proteins on another member of the protease-activated receptor family (15) we hypothesized that endogenous PAR1 signaling could be affected by SR12 expression. PAR1 signaling involves Gα_q/11_ activation, triggering phospholipase C (PLC) and the generation of diacylglycerol (DAG). To assess changes in DAG production in cells endogenously expressing PAR1 after SR12 transfection, a BRET-based biosensor was employed to detect this molecule, as outlined in the methods and illustrated in Figure 7A.

**Figure 7.**
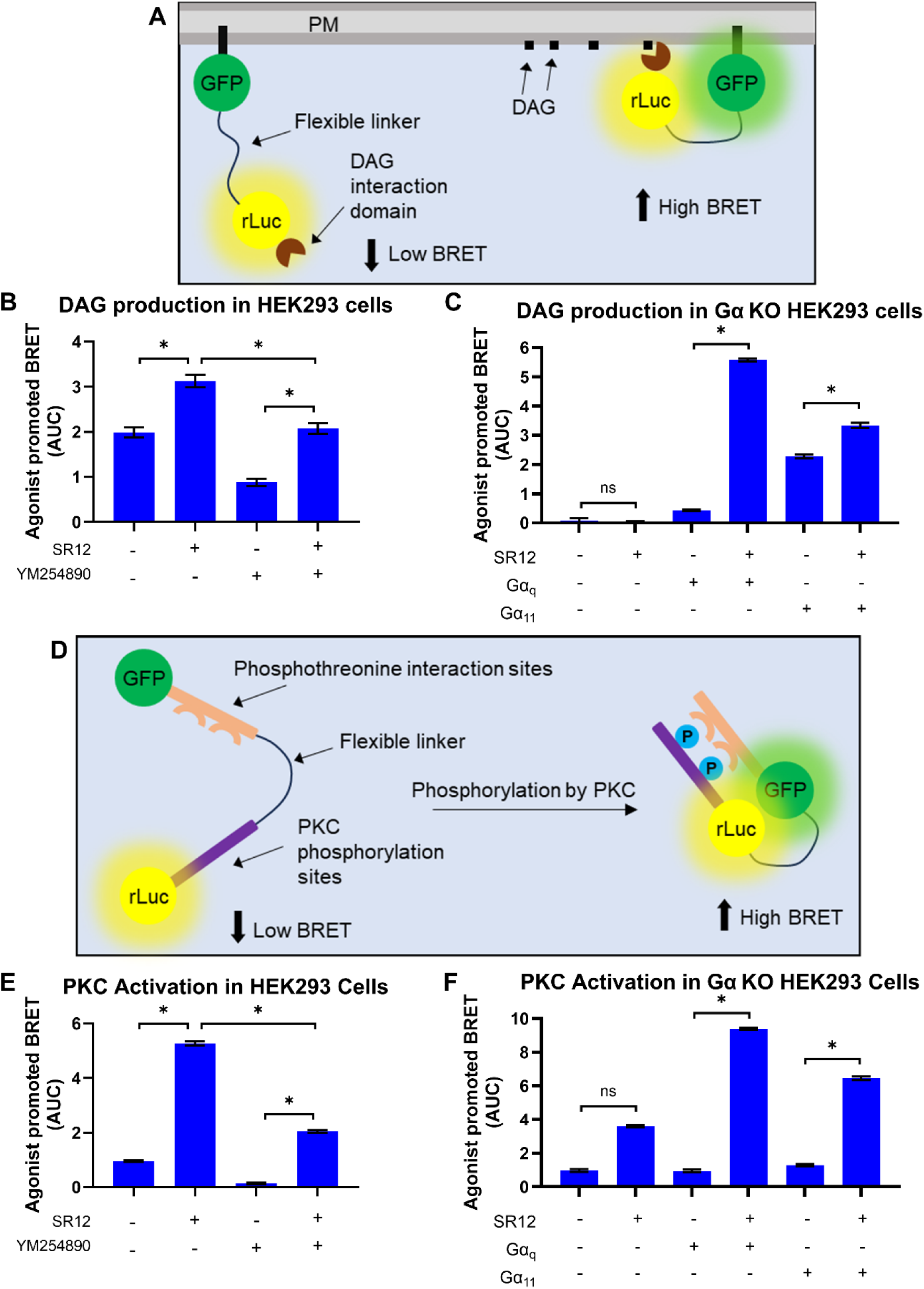
DAG formation and PKC activation measurements. (**A**) Functioning model of DAG biosensor. (**B**) HEK293 cells were co-transfected with SR12 or mock plasmid, and DAG biosensor. Cells were treated with thrombin (2U/mL) or vehicle. Gα_q/11_ inhibitor (YM-254890) was added 30 minutes before reading the experiment. (**C**) HEK293 G_α_ KO cells were co-transfected with SR12 or mock plasmid and DAG biosensor and treated with thrombin (2U/mL) or vehicle. G_α_ protein rescue experiment was performed by adding Gα_q_ or Gα_11_ to the transfection mix. (**D**) Functioning model of PKC biosensor. (E) HEK293 cells were co-transfected with SR12 or a mock plasmid and PKC biosensor. Cells were treated with thrombin (2U/mL) or vehicle. Gα_q/11_ inhibitor (YM-254890) was added 30 minutes before reading the experiment. (**F**) HEK293 Gα KO cells were co-transfected with SR12 or a mock plasmid and PKC biosensor and treated with thrombin (2U/mL) or vehicle. Gα protein rescue experiment was performed by adding Gα_q_ or Gα_11_ to the transfection mix. BRET values were calculated by dividing the intensity of light emitted by rGFP (515 nm) by the intensity of light emitted by rLuc2 (400 nm) during a 5-minute kinetic loop with successive readings. Area under the curve was then calculated by setting the baseline (before compound addition) to 0. Results were normalized by subtracting the area under the curve of vehicle-treated cells from the area under the curve of thrombin-treated cells (mean ± SEM, n=3). * Statistically significant by two-way ANOVA with Šidák post-test, p<0,05.

As shown in Figure 7B, SR12 expression significantly potentiated DAG production after stimulation by thrombin. This response was blocked by pre-treatment with the Gα_q/11_ family-selective protein inhibitor, YM-254890. The DAG response observed in the condition without SR12 expression likely results from the endogenously expressed thrombin receptor in HEK293 cells, which have been shown to express PAR1, PAR2, and PAR4 (32).

To further validate Gα_q/11_ family activation, knockout cells for this family of G proteins were employed, with the subsequent rescue of each protein expression. As expected, no DAG formation was observed in the Gα_q/11_ knockout cell line upon thrombin stimulation while the response was rescued upon re-expression of Gα_q_ and Gα_11_, and resulted in a significantly greater increase in DAG formation in SR12- transfected cells compared to mock conditions (Figure 7C). The lack of a complete inhibition of DAG production after YM-254890 treatment is likely due to the Gβγ signaling released from Gα_i/o_, strongly activated by protease activated receptors (33); this observation is further supported by the complete absence of signalling in GαKO cells (Figure 7C).

To assess whether the potentiation of DAG formation could also be detected in downstream signaling events, we examined the capacity of thrombin to stimulate protein kinase C (PKC) in the presence and absence of SR12 expression using a BRET-based PKC sensor (34) as described in the methods and illustrated in Figure 7D. Consistent with expectations, PKC activation in response to thrombin stimulation was higher in SR12-transfected cells compared to cells not expressing SR12, and the signal was largely blocked upon treatment with YM-254890 (Figure 7E). The PKC activation signal also was attenuated in knockout cells for the Gα_q/11_ family but restored upon supplementation with Gα_q_ and Gα_11_ (Figure 7F). Interestingly, the SR12-potentiation observed for the PKC response was larger than that observed for DAG response, most-likely resulting from an amplification of the signal. In addition, we also detected an SR12 mediated potentiation of the cytoplasmatic [Ca^2+^] rise using Obelin-based luminescent biosensor (Supplementary Figure S4). Of note, based on AF models, it is unlikely that SR12 couples directly to the G protein found in HEK293T cells due to a steric clash between the G protein and the SR12 Helix 8 (Supplementary figure S5) and the lack of canonical switch motifs (PIF, NPXXY, DRY) in its primary sequence. Taken together, these results indicate that SR12 can potentiate the activation of downstream effectors of Gα_q/11_/PLC/DAG such as PKC in response to thrombin, indicating that this 7 TM-protein have a clear cell signaling impact, most likely due to its action on PAR1 trafficking leading to increased cell surface and organelle expression.

Since HEK293 cells have significant endogenous expression of PAR1, we investigated whether SR12 could affect signaling in PAR knockout cells. Parental cells that endogenously express PAR showed increased DAG production upon SR12 transfection (Figure 8A). On the other hand, in the absence of endogenous PAR, SR12 expression did not promote thrombin-promoted DAG production, indicating that SR12 cannot act as a thrombin-activated receptor (Figure 8A). Over-expression of PAR1 in the PAR-KO cells restored a robust thrombin-stimulated DAG response (Figure 8B). These results strengthen the hypothesis that SR12 promotes PAR1 signaling by increasing the low expression levels of the endogenous receptor.

**Figure 8.**
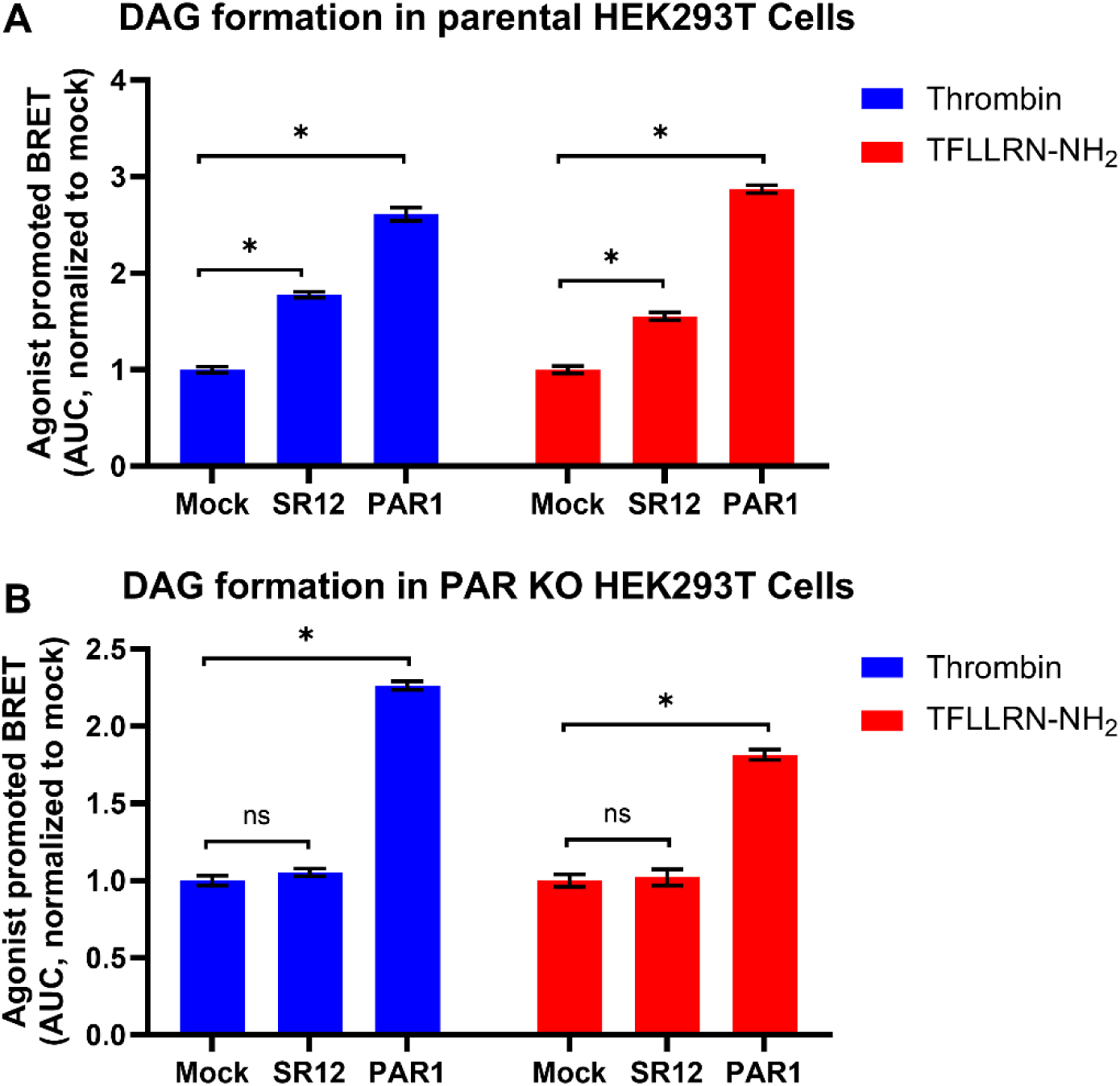
SR12 transfection in PAR1 knockout cells. (**A**) HEK293 parental cells were co-transfected with SR12 or PAR1 and DAG biosensor and then treated with 2U/mL thrombin or 10µM of the PAR1 tethered ligand TFLLR-NH_2_. (**B**) HEK293 PAR KO cells were co-transfected with SR12 or PAR1 and DAG biosensor and then treated with 2U/mL thrombin or 10µM of the PAR1 tethered ligand TFLLR-NH_2_. BRET values were calculated by dividing the intensity of light emitted by rGFP (515 nm) by the intensity of light emitted by rLuc2 (400 nm) during a 5 minute kinetic loop with successive readings. Area under the curve was then calculated by setting the baseline (before compound addition) to 0. Results were normalized as fold-change relative to mock transfected cells (mean ± SEM, n=3). * Statistically significant by two-way ANOVA with Šidák post-test, p<0,05.

Given that SR12 potentiated the response of an endogenously expressed PAR, we wondered whether it might be a more general phenomenon that potentiates other GPCR responses. To test this hypothesis, we assessed changes in response to carbamoylcholine, an agonist of endogenously expressed muscarinic receptors (Figure S6). Using DAG and PKC biosensors, we found that SR12-transfected cells showed a greater response to Carbamoyl Choline than cells without SR12 expression. This result indicates that SR12 can increase signaling by other endogenous GPCRs, not exclusively PAR1.

## Discussion

Our data suggest that SR12 is a member of the GOLD-domain 7TM protein family that can regulate the subcellular distribution of GPCRs when expressed in a heterologous system. This classification is supported by: (1) three-dimensional structure prediction via AlphaFold, molecular dynamics simulations and alignment with known members of the GOLD-domain 7TM family, including GPR107, GPR108, GPR180, and TMEM87A; (2) the observed increase in the distribution of at least two GPCRs across subcellular compartments, including the secretory pathway, plasma membrane, and endosomes; and (3) the potentiation of signaling mediated by thrombin- and carbamylcholine-activated endogenously expressed GPCRs.

Based on our findings that SR12 may assist in GPCR trafficking in heterologous systems, we propose two potential roles for this protein: (1) acting as a chaperone for parasite-encoded GPCR-like proteins, such as SR1, SR10, and SR25, to ensure their proper cellular distribution and function; or (2) interfering with the distribution of GPCRs within the infected cell, thereby sensitizing the parasite to host-derived signals, such as changes in Ca²⁺ or cAMP concentrations.

Protein synthesis and trafficking are complex processes during *Plasmodium*’s life cycle in the human host, due to the unique environment that proteins must traverse, which involves multiple lipid membranes, including the plasma membrane, parasitophorous vacuole membrane, and erythrocyte plasma membrane. Moreover, proteins may localize to the erythrocyte cytoplasm or to organelles that are either specific to *Plasmodium* or functionally distinct from their human counterparts, such as the mitochondrion, digestive vacuole, or apicoplast. Thus, *Plasmodium* possesses specialized mechanisms and proteins responsible for assisting in the trafficking and localization of other proteins. Heat Shock Proteins (HSPs) are the primary class of proteins responsible for maintaining proteostasis throughout the parasite life cycle, exhibiting superior chaperone activity compared with their human counterparts. Within this family, HSP70 stands out for its roles in aggregation suppression (35, 36); ER protein folding/export (37, 38); mitochondrial folding of asparagine-rich proteins (39, 40); and protein transport to the host cell cytoplasm (38). While HSPs functions are well-characterized in soluble protein folding and trafficking, little is known about how *Plasmodium* cells manage these challenges for complex multi-transmembrane domain proteins, and SR12 could take part in this process.

GPCRs exhibit complex topological structures and are inherently prone to folding challenges, which complicate their expression, trafficking, and stabilization at the plasma membrane. To circumvent these issues, cells rely on GPCRs’ partner mechanisms that assist in folding, stabilization, and localization, such as heterodimerization of CaSR and mGlu5 or GABAb2 (41, 42); heterodimerization of GABAb2 and GABAb1; homodimerization of β2AR (43); and associations between RAMPs and various GPCRs. In *Plasmodium*, some GPCR-like proteins may exploit dimerization with SR12 for trafficking or even functional modulation, including: SR25, a monovalent cation sensor linked to antimalarial drug resistance (5, 44); the ER-resident SR10, associated with cellular differentiation (45); and SR1, whose function remains unknown. Additionally, we cannot exclude that SR12 can be used by the parasite to hijack host-cell signalling components, leveraging GPCRs and effector proteins already present in the host cell to promote parasite development. This mechanism is very well known in viral infections, but also reported in bacterial and eukaryotic parasites, such as: *Staphylococcus aureus* toxins interactions with chemokine receptors ACKR1, CXCR1, CXCR2, CCR2, and CCR5 (46, 47); *P*. *gingivalis* proteases altering PAR2 dependent Ca^2+^ signalling (48); *N. meningitidis* mechanical activation of β2AR (49); and Toxoplasma gondii using CCR7/CCL19-mediated chemotaxis to increase migratory proprieties of the host cell (50, 51). Interestingly, it was demonstrated in *P. falciparum* infection that blocking either β2AR or Gαs signalling caused a decrease in both *in vitro* and *in vivo* parasite infection (52), confirming the importance of GPCRs and GPCR signalling in the parasite’s lifecycle. In this context, SR12 could act in association with endogenous GPCRs changing subcellular localization and/or increasing signalling during infection of hepatocytes and erythrocytes in order to promote parasite survival.

All homologs of SR12 represented in Figure 1 are members of the GOST family proteins. Although these proteins are recognized for their role in cargo transport along the Gogi network, their target cargoes and the potential specificity of these proteins for specific cargos remain poorly understood. One example of the role of GOST family members in the secretory pathway is the transport of lipidated Wnt by Wntless, also known as GPR177, between internal membranes and the cell surface (30, 53, 54). In addition, TMEM145, which is another member of GOST family and a very close homolog of SR12 (figure 1C), has recently been shown to be involved in the transport and secretion of several proteins responsible for the formation and integrity of stereo ciliary structures in outer hair cells of cochlea, within inner ear (55).

Although none of the GOST family proteins have been shown to bind to GPCRs as their target cargo, TMED2 or TMED10 proteins which are integral proteins with a GOLD domain, have been shown to interact with and modulate the trafficking of several GPCRs such as Smoothened (SMO), calcium sensing receptor (CaSR), Protease-activated receptor-2 (PAR-2), μ-opioid receptor-1B (MOR1B) and several purinergic GPCRs such as P2Y_1_, P2Y_2,_ P2Y_4_ and P2Y_11_ (56–59). In case of PAR-2, the GOLD domain of TMED2 interacts with the extracellular loop 2 (ECL-2) of PAR-2 and deletion of GOLD domain in TMED2 results in increased cell-surface expression of PAR-2 (58) indicating that its effect on GPCR traficking is distinct from the one observed here for SR12.

The orphan GPCR GPR107, which shares a three-dimensional structural similarity with SR12, is predominantly expressed in the Golgi apparatus (28). Its role in trafficking has recently been reported: its deficiency inhibits endocytosis of angiotensin receptor 1 (AT1R), thereby altering angiotensin-mediated calcium flux (12). GPR107’s interaction with clathrin and vacuolar protein sorting ortholog 35 (VPS35) has also been demonstrated, directly linking this receptor to cargo transport and membrane receptor trafficking (27). Another analogous GPCR, GPR108, is also localized to the Golgi apparatus (13) and participates in adeno-associated virus transduction (9, 10). Recently, GPR180, the human protein with the highest similarity to SR12, was shown to localize to intra-cellular organnelles’ membranes (11). In this context, it can be proposed that SR12 facilitates the transport of other 7TM proteins to overcome barriers in *Plasmodium* protein trafficking.

The structural predictions and functional similarities indicate that SR12 belongs to the GOST protein family. This family comprises several orphan GPCRs and other transmembrane proteins involved in intracellular transport. SR12’s ability to enhance the abundance of GPCRs at the plasma membrane, endosomes, and the Golgi apparatus aligns with the known functions of other members of this family. These findings provide insights into the potential role of SR12 in host-parasite interaction and shed light on the function of human GOST proteins.

## Study limitations

The major limitation of the data presented herein is that we used heterologous expression systems in mammalian cells to study the function of SR12. Although this approach provided insights into SR12 interactions in mammalian cells and the overall functions of GOLD domain proteins, it is not possible to confirm their function in *Plasmodium* cells, as they do not express canonical GPCRs. Further experiments will explore the role of this protein in *Plasmodium* infected erythrocytes and hepatocytes. There is also a gap in structural knowledge of *Plasmodium* proteins due to lack of CryoEM, NMR and x-ray resolved structures, preventing confirmation of the AF modeled structures with experimental structural data. The predictions, together with MD simulations and comparison with GPR180 revealed the presence and the function of a GOLD domain involved in transport that was supported by our cell-based assays, but the role of the seven transmembrane domain as well as the long unstructured N-terminal part of the protein remain to be elucidated.

## Materials and Methods

### Cell culture and transfection

Human Embryonic Kidney 293 SL (HEK293SL) cells were cultivated in T150 flasks with Dulbecco’s Modified Eagle Media (DMEM) supplemented with 10% Newborn Calf Serum (NCS), 100 units/mL penicillin and 100 µg/mL streptomycin. The flasks were kept at 37°C/0.5% CO2 incubator. Human Embryonic Kidney 293T parental cells and knockout cells for PAR receptors or Gα_q/11_ proteins (HEK293T parental, HEK293T PAR-KO and HEK293 Gα_q/11_-KO) were cultivated in T150 flasks with Dulbecco’s Modified Eagle Media (DMEM) supplemented with 10% Fetal Bovine Serum (FBS), 100 units/mL penicillin and 100 µg/mL streptomycin. The flasks were kept at 37°C/0.5% CO2 incubator.

HEK293SL cells were co-transfected with pCDNA3.1(-) plasmid containing the codon optimized versions of SR12 (PF3D7_0422800) and plasmids containing components of the biosensor used. Transfection of cells were done using Branched Polyethylenimine (PEI, Millipore-Sigma) in a proportion of 3 µg per 1 µg of DNA directly in culture media containing 350.000 cells/mL and seeded in a 96 well plate using 100µL of DNA-Cell mix aiming a final concentration of 300.000 cells/mL. For HEK293T cells, transfection was done using 500.000 cells/mL aiming at a final concentration of 41.000 cells/mL in a 96 well plate. Cells were then kept for 48 hours at 37°C before luminescence or BRET experiments were performed.

### Calcium mobilization measurement

Calcium mobilization measurements were performed by transfecting HEK293SL cells with a plasmid containing an Obelin biosensor. Obelin is an Aequorin analog that consists of a single 22kDa polypeptide chain with a high affinity for 2-hydroperoxycoelenterazine. Obelin can consume its substrate to produce blue light spontaneously, but in the presence of free Ca^2+^, the intensity of the light emission is one million times higher (Illarionov et al. 2000; Gealageas et al. 2014). Cells were washed in PBS buffer (137 mM NaCl, 2,7 mM KCl, 4,3 mM Na_2_HPO_4_, 1,4 mM NaH_2_PO_4_, pH 7.4), incubated for 2 hours with 100 nM Coelenterazine Cp and washed before initiating measurements using a spectrophotometer (SpectraMax M, Molecular Devices) equipped with injectors allowing the detection of luminescence in a kinetic mode.

### Bioluminescence Resonance Energy Transfer (BRET) assay

Before reading, cells were washed in PBS buffer (137 mM NaCl, 2,7 mM KCl, 4,3 mM Na_2_HPO_4_, 1,4 mM NaH_2_PO_4_, pH 7.4) and incubated 15 minutes with 100 nM bis-desoxi-coelenterazine (DeepBlueC^TM^) before initiating BRET measurements using a spectrophotometer (Mithras LB 940, Berthold Tech.) equipped with filters allowing the detection of luminescence at different wavelengths simultaneously. BRET values in the presence of different compounds were monitored in a kinetic mode. The time-dependent differences in BRET values were quantified and used as a readout for the activation of the pathways monitored (see the description of the biosensors used below). BRET values were determined by rationing the luminescence emitted at 510 nM (acceptor) over the luminescence emitted at 395 nM (donor).

The DAG biosensor is a unimolecular BRET-based sensor used to detect the formation of diacylglycerol in the plasma membrane. It is composed of a protein chimera containing a domain that attaches the biosensor to the membrane through myristoylation/palmitoylation, a GFP (acceptor), a long flexible linker, rLuc (donor) and a DAG interaction domain c1b from PKCδ (34). The PKC biosensor is a construct of an rLuc2 conjugated to two PKC phosphorylation sites (TLKI and TLKD) linked by a flexible linker to phosphotreonine interaction domains (FHA1 and FHA2) and a GFP (Namkung et al. 2018a). The subcellular localization using BRET was done transfecting C-terminally rLuc2 tagged versions of each receptor together with an rGFP fused to a peptide targeted to different subcellular compartments: rGFP-CAAX targeting plasma membrane; rGFP-FYVE targeting early endosomes (60); rGFP-Giantin for Golgi apparatus; rGFP-Bcl-xL targeting mitochondria (61).

### Sub cellular localization of SR12 by fluorescence microscopy

A codon-optimized variant of monomeric Kusabira-Orange2 (mKO2, (62)) was synthesized and subsequently subcloned into the multicloning site of a mammalian expression vector, pSF-CMV-AMP, using NcoI and XbaI sites. To achieve plasma membrane, early endosomes, Golgi apparatus and ER localization, the K-Ras-derived CAAX motif, FYVE domain of endofin (60), Golgi targeting domain of giantin (61) and the C-terminus of SecG (beta-subunit of Sec61) were, respectively, fused to the C-terminus of mKO2 using Gibson assembly (Supplementary Figure S7).

After reaching 85% confluency, Human Bone Osteosarcoma Epithelial cells (U2OS) were resuspended in DMEM supplemented with 10% FBS and 1% of penicillin-Streptomycin. Transfection was performed using 1μg of total DNA (600ng of GFP-SR12, 100ng of mKO2-Sec61b, 200ng of mKO2-CAAX, mKO2-FYVE and mKO2-Giantin and adjusted amounts of salmon sperm DNA) using X-tremeGENE 9 DNA (Millipore Sigma #6365787001) diluted in Opti-MEM I 1x (Thermo Fisher # 31985070) as a transfection agent, following the 3:1 ratio of X-tremeGENE:DNA. Cells were added to the transfection mix (DNA+X-tremeGENE) and immediately seeded into a 35mm tissue culture dish with glass bottom at a concentration of 2.5 x 105 cells per well and maintained in culture for 48 hours. Images were acquired using the LSM880 confocal microscope (Zeiss) with 40x objective.

### Structural prediction and Structural homology search

We performed structural predictions for full-length SR12 using Alphafold2colab and standard input parameters (63). The structure with the highest pLDDT score was selected, and the structure covering residues 94-470 was used for MD simulation as well finding homolog proteins. To further probe the functional aspects of the predicted model and finding close structural homologs, we queried the Foldseek web server (https://search.foldseek.com) in 3Di/AA mode against two databases: PDB100 (non-redundant experimentally determined PDB structures) and the Swiss-Prot subset of AlphaFold-DB (https://alphafold.ebi.ac.uk/download#swissprot-section). In 3Di/AA mode, each structure is encoded as a sequence over a 20-state 3Di alphabet (capturing local 3D residue-residue interactions), and alignments are scored using both 3Di and amino-acid substitution information to increase sensitivity (64). We assessed significance of hits with Foldseek’s calibrated E-values, which estimate the expected number of chance matches per query with score at least as high as the observed alignment (64). We retained hits with E ≤ 1×10^-3^, a threshold that corresponds to ≤0.001 expected random matches per query (≈≤1 false positive per 1,000 queries) and provides a practical balance between sensitivity and specificity. Under this threshold, 33/520 hits were kept for downstream analyses.

### System preparation and molecular dynamics simulation

To insert the receptor into 1-palmitoyl-2-oleoyl-sn-glycerol-3-phosphatidylcholine (POPC) bilayer model, we utilized the CHAMM-GUI membrane builder server (65). The system was neutralized with NaCl counter-ions to a concentration of 0.15 M. MD simulations were conducted using GROMACS 2021.2, employing the CHARMM36m force field for the protein and lipids (66, 67). The CHARMM TIP3P water model served as an explicit solvent. The simulation system underwent consecutive energy minimization, equilibration, and production runs, utilizing scripts generated by CHARMM-GUI. The systems were energy-minimized with steepest descent algorithms, followed by six-step equilibration runs. Restraint forces were progressively reduced during the equilibration process. An unrestrained NPT production run of 1000 ns was performed with periodic boundary conditions along all three orthonormal directions. The particle mesh Ewald (PME) method calculated long-range Coulombic interactions, with a cut-off distance of 1.2 nm. A force-switch function was applied for Lennard-Jones interactions, featuring a cut-off of 1.0 nm. The temperature was maintained at 310 K using the Nose-Hoover thermostat, and the system pressure was kept at 1 bar with the Parrinello-Rahman barostat. The covalent bonds between hydrogen and other heavy atoms were constrained by LINCS algorithm, permitting a simulation time-step of 2 fs.

### Trajectory analysis and visualization

Production run trajectories were analyzed using GROMACS tools. All molecular graphics work and figures were generated using UCSF Chimera and VMD software (ref). PCA analysis was performed using “gmx covar” tool of GROMSCS to yield the eigenvalues and eigenvectors by calculating and diagonalizing the covariance matrix, whereas the “gmx anaeig” tool was used to analyze the eigenvectors. The eigenvalues were obtained by the diagonalization of the covariance matrix of the Cα atomic fluctuations. To represent the movement directions captured by the eigenvectors, the porcupine plot was generated using 30 extreme projections on principal components PC1, as the input for the Prody plugin of VMD software (68). To measure and represent protein cavities, the first and last frames of simulation were analyzed by CASTp 3.0 server (http://sts.bioe.uic.edu/castp/index.html?3trg). The volume of the main protein pocket was evaluated by POcket Volume Measurer (POVME) version 2.0, using the defaults settings (69). As input for the analysis, 100 evenly distributed structures were selected from the whole simulation. The electrostatic surface potential of protein was calculated using APBS-PDB2PQR server (https://server.poissonboltzmann.org/).

## Acknowledgments

This work was supported by the Fundação de Amparo à Pesquisa do Estado de São Paulo (FAPESP) grants to CRSG (23/07656-0), PHSP (2017/16307-9 and 2014/14347-5), and CLK (2016/09185-1), by a Foundation grant of the Canadian Institute for Health Research (FDN148431) and a project grant (PJT-183158) to MB. This research was enabled in part by support provided by the Calcul Québec (https://www.calculquebec.ca) and the Digital Research Alliance of Canada (https://alliancecan.ca).

---

**Figure S1.**
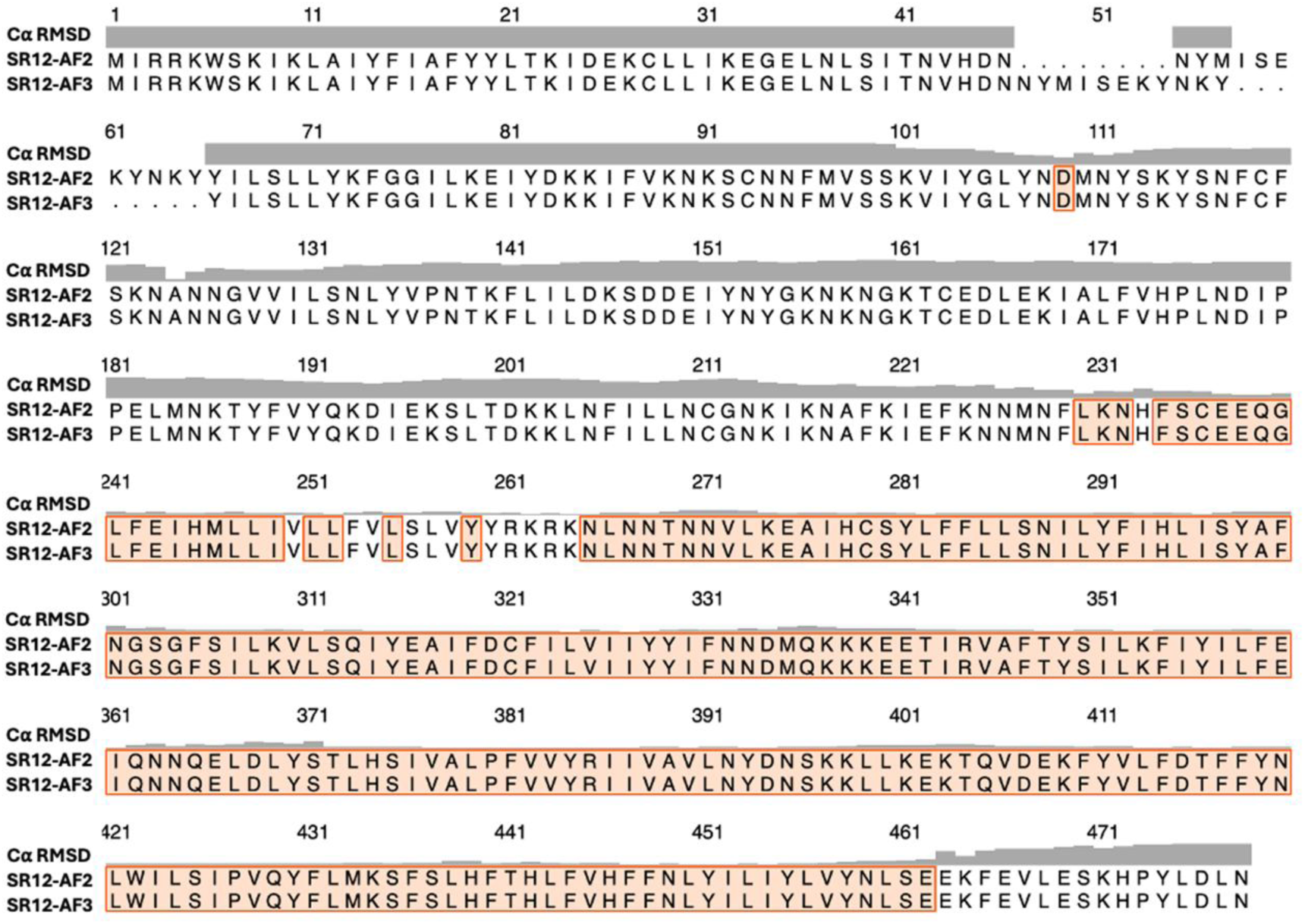
The pairwise sequence alignment of AF2 and AF3 models representing the residues that are overlaid from each model and the RMSD of their C⍺ atoms. The residues pairs with lowest RMSD are highlighted.

**Figure 2S.**
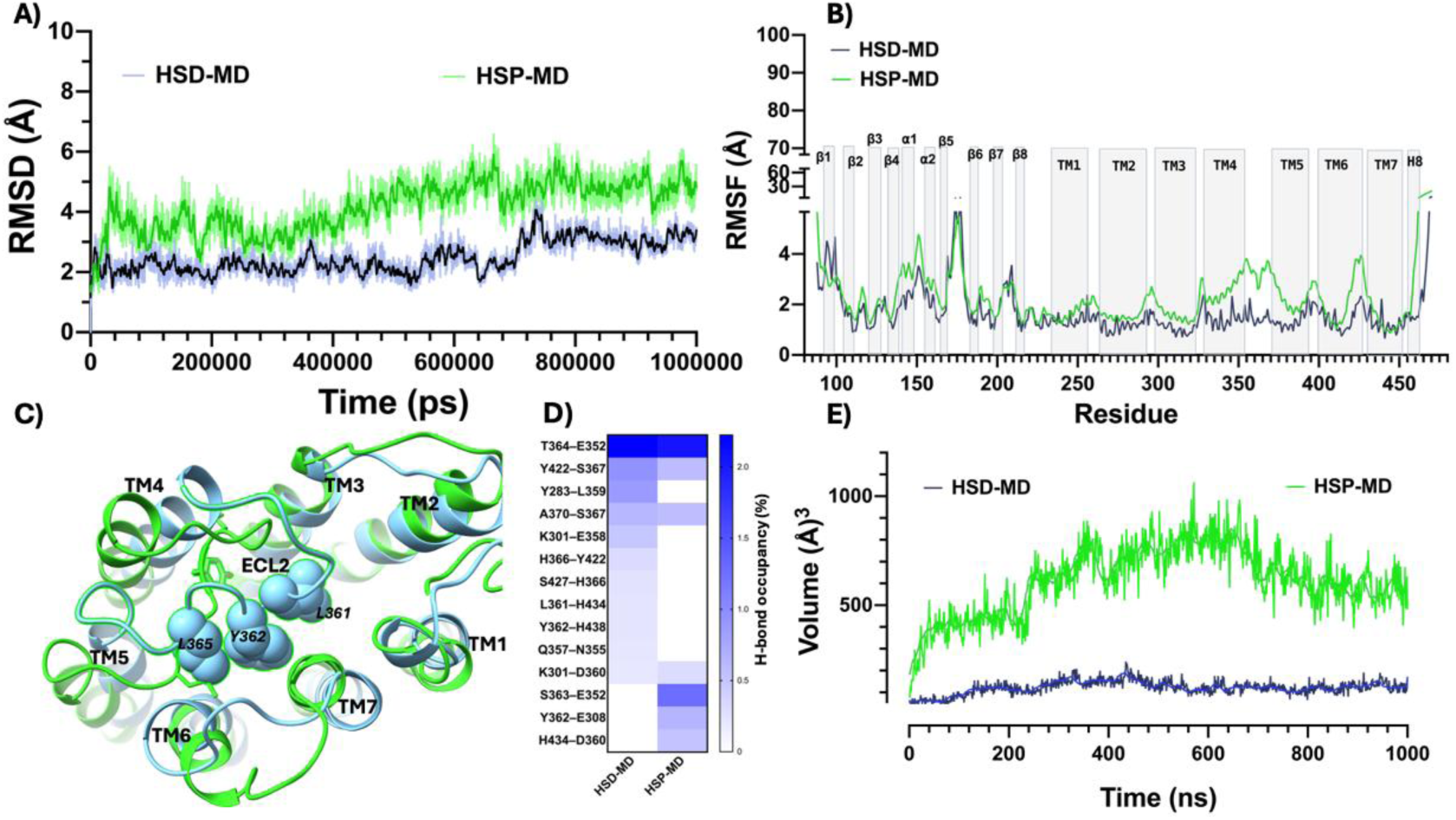
Protonation of extracellularly exposed histidines increases conformational drift and expands the SR12 extracellular pocket. (A) Time evolution of backbone RMSD for simulations with neutral histidines (HSD; blue) and protonated histidines (HSP; green). (B) Per-residue RMSF for HSD (blue) and HSP (green), highlighting increased mobility in extracellular loops. (C) Representative snapshots from HSD (blue) and HSP (greenn) trajectories illustrating ECL2 reorientation. (D) Protonation-dependent remodeling of the ECL2 hydrogen-bond network, summarized as changes in hydrogen-bond occupancy across the trajectories. (E) Extracellular pocket volume as a function of time, showing pocket expansion in HSP (green) relative to HSD (blue).

**Figure S3.**
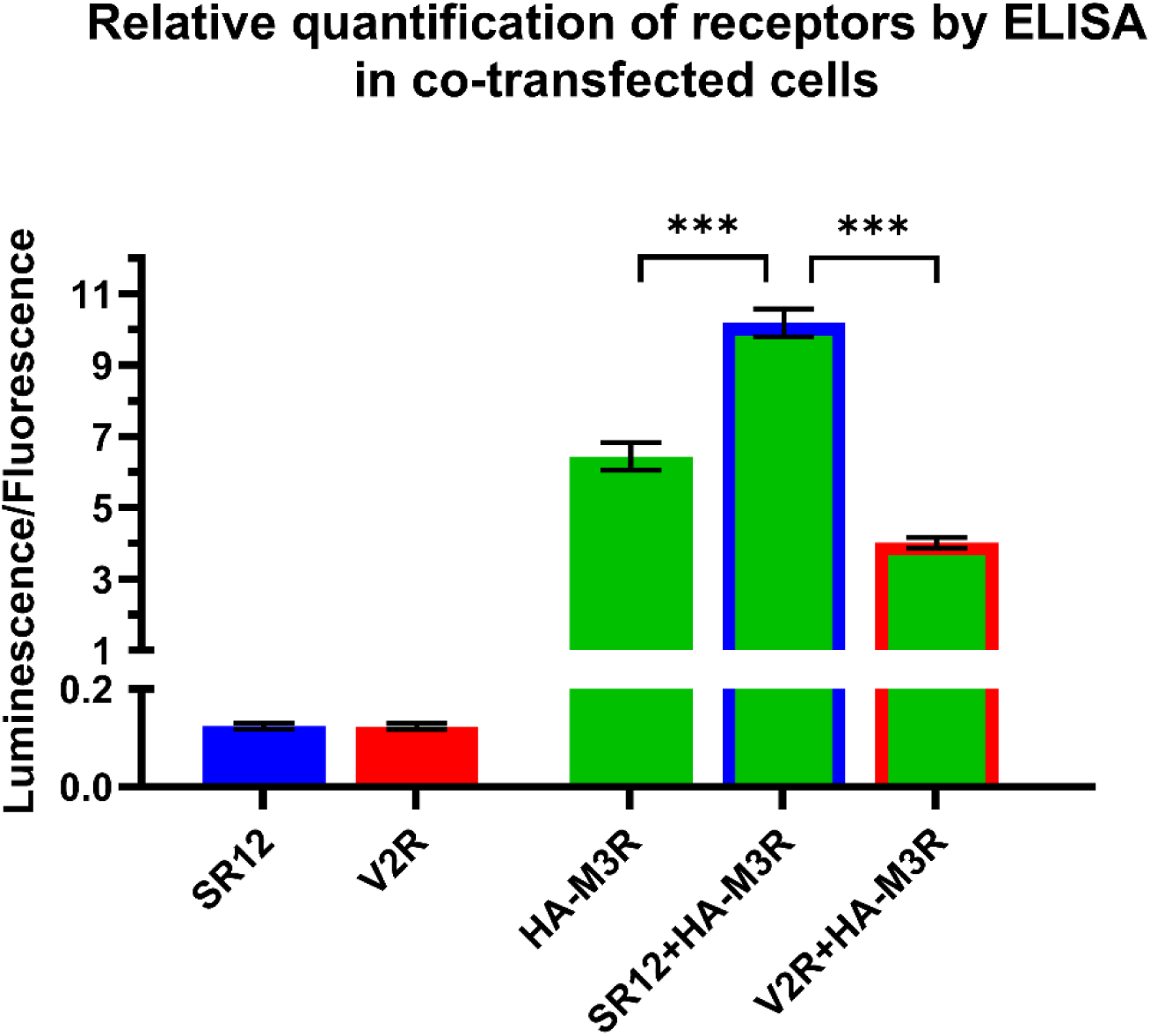
Plasma membrane expression of SR12 and selected GPCRs by ELISA. HEK293SL cells were transfected using 200ng of each receptor and detected on the membrane surface by ELISA using anti- HA antibody. SR12 and V2R untagged versions were used, so no signal was expected for these receptors. M3R HA-tagged versions was used, which showed a positive luminescence increase. When SR12 is cotransfected with M3R-HA, the signal is boosted by 58%. On the other hand, when V2R is co-transfected with M3R-HA, the expression decreases 38%, as expected due to competition between plasmid expression. * Statistically significant by two-way ANOVA with post-test T.

**Figure S4.**
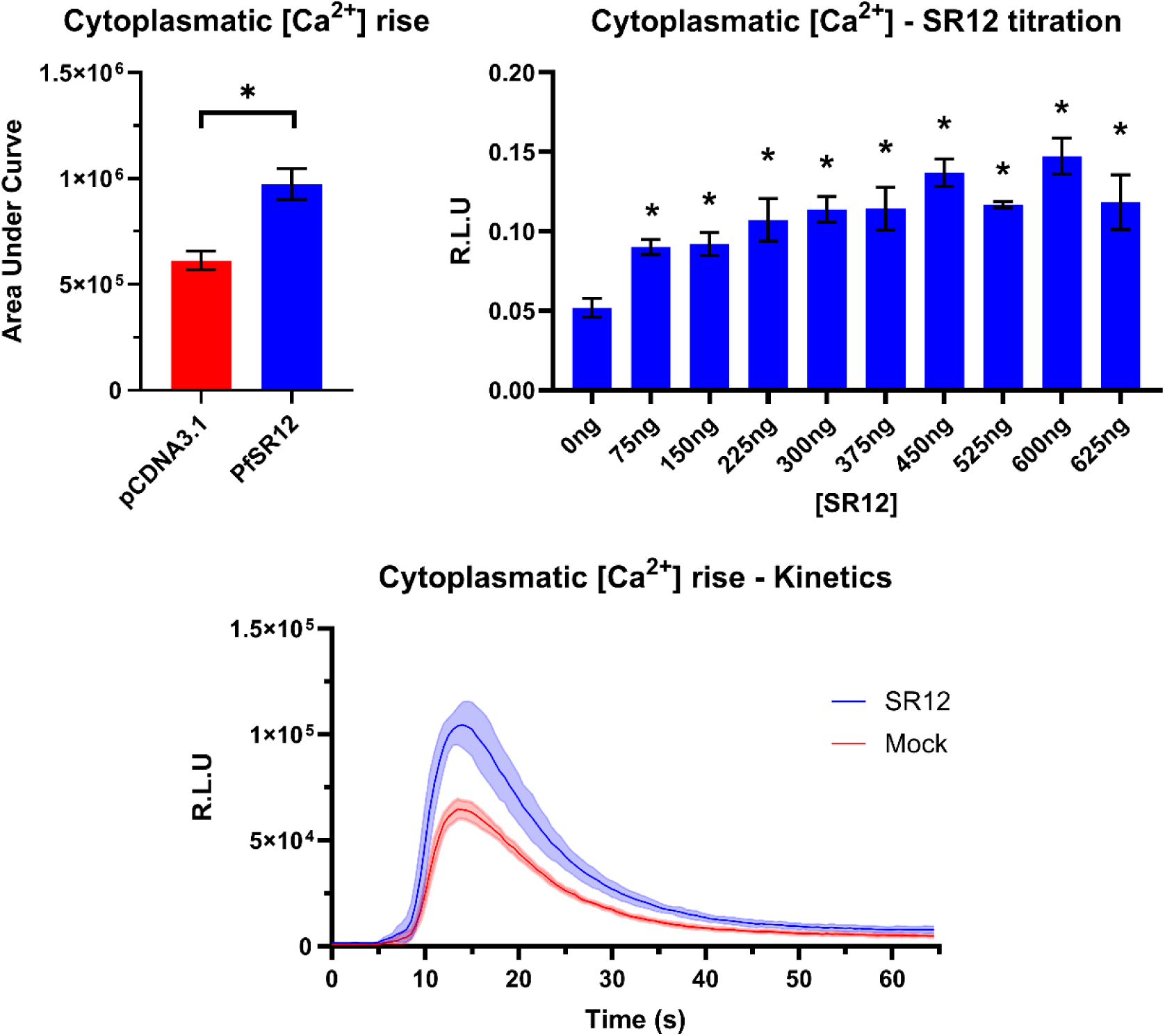
Cytosolic Ca2+ response after dose-dependent thrombin treatment in HEK293SL cells transfected with SR12. HEK293SL cells were co-transfected with Obelin/Ca2+ biosensor and SR12. 2U/mL thrombin induces Ca2+ rise 1.59 times higher in SR12 transfected cells when compared to mock cells (A). This Ca2+ rise is dependent of the amount of SR12 transfected (B). (C) shows a typical kinetic pattern for intracellular calcium release and subsequent uptake. * Statistically significant by Student’s T test for two means (A). For figure B, statistically significant by two-way ANOVA with post-test T and P <0.05 (*) or P <0.001 (***).

**Figure S5.**
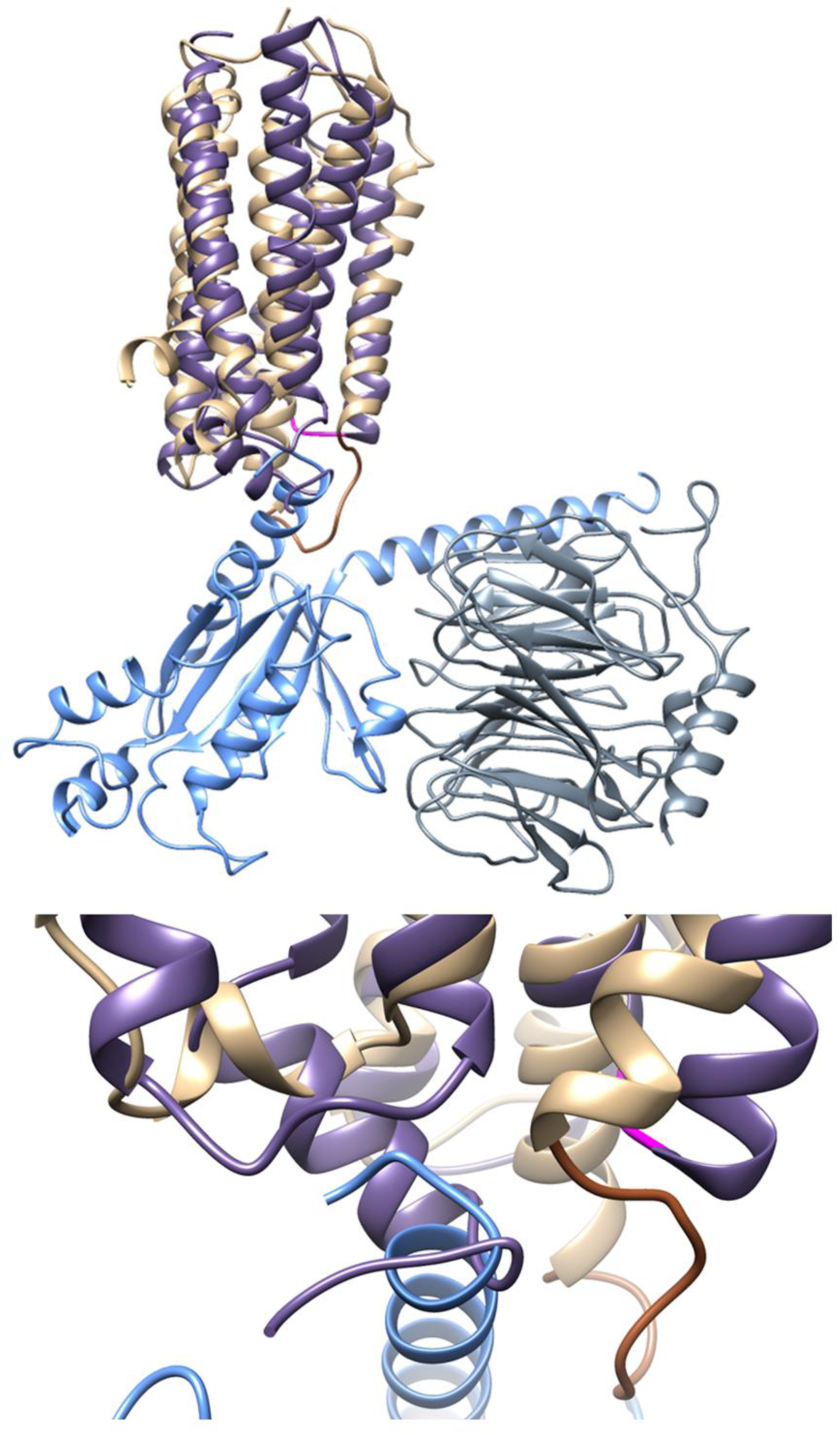
Overlay of SR12 structure on the active form of mGlu2 structure in complex with G⍺i-Gβɣ. The steric clash between the H8 of SR12 with the G-protein as well as the unconventional length and orientation of ICL2 are highlighted.

**Figure S6.**
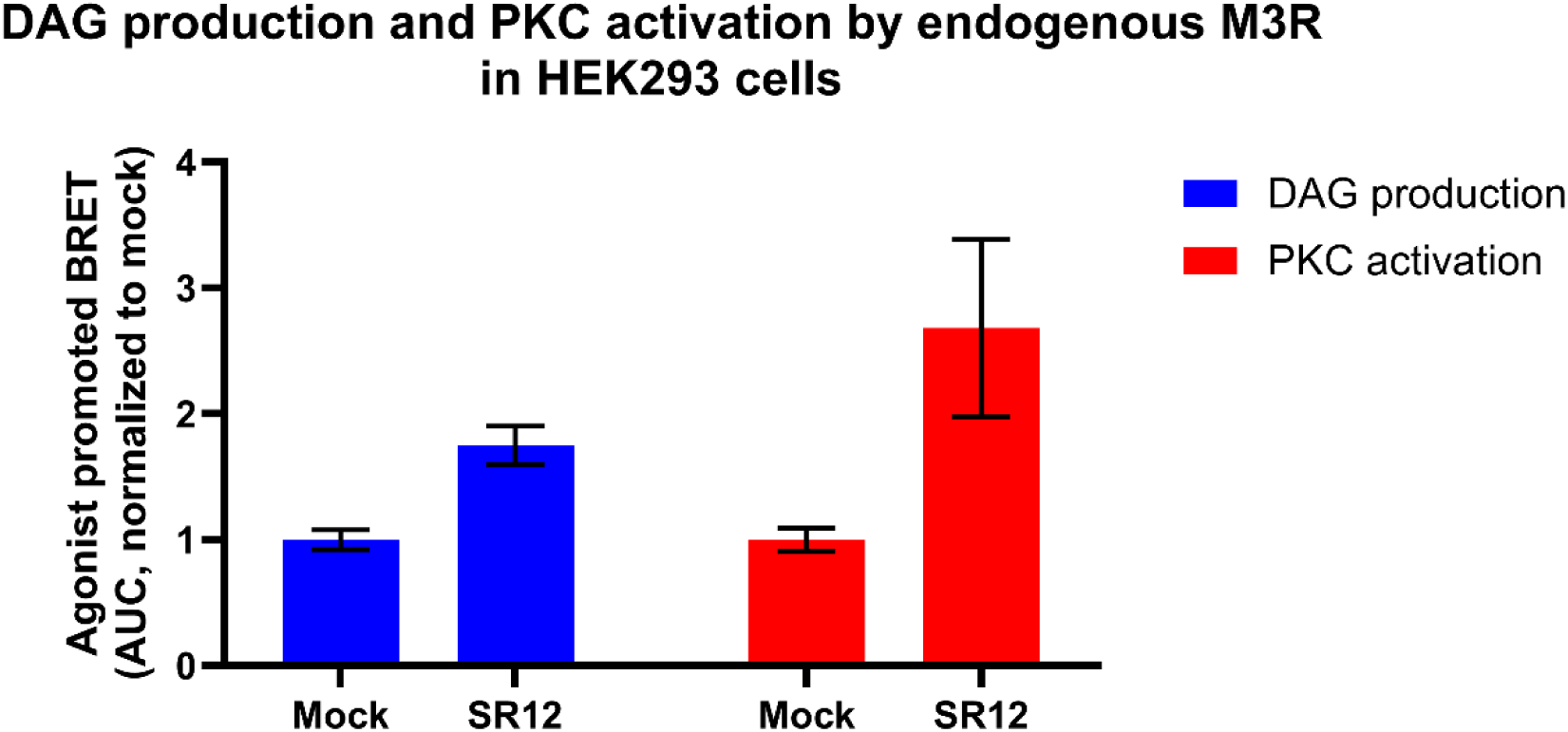
SR12 transfection in PAR1 knockout cells. (**A**) HEK293 parental cells were co-transfected with SR12 or PAR1 and DAG biosensor and then treated with 2U/mL thrombin or 10µM of the PAR1 tethered ligand TFLLR-NH_2_. (**B**) HEK293 PAR KO cells were co-transfected with SR12 or PAR1 and DAG biosensor and then treated with 2U/mL thrombin or 10µM of the PAR1 tethered ligand TFLLR-NH_2_. BRET values were calculated by dividing the intensity of light emitted by rGFP (515 nm) by the intensity of light emitted by rLuc2 (400 nm) during a 5 minute kinetic loop with successive readings. Area under the curve was then calculated by setting the baseline (before compound addition) to 0. Results were normalized as fold-change relative to mock transfected cells (mean ± SEM, n=3). * Statistically significant by two-way ANOVA with Šidák post-test, p<0,05.

**Figure S7.**
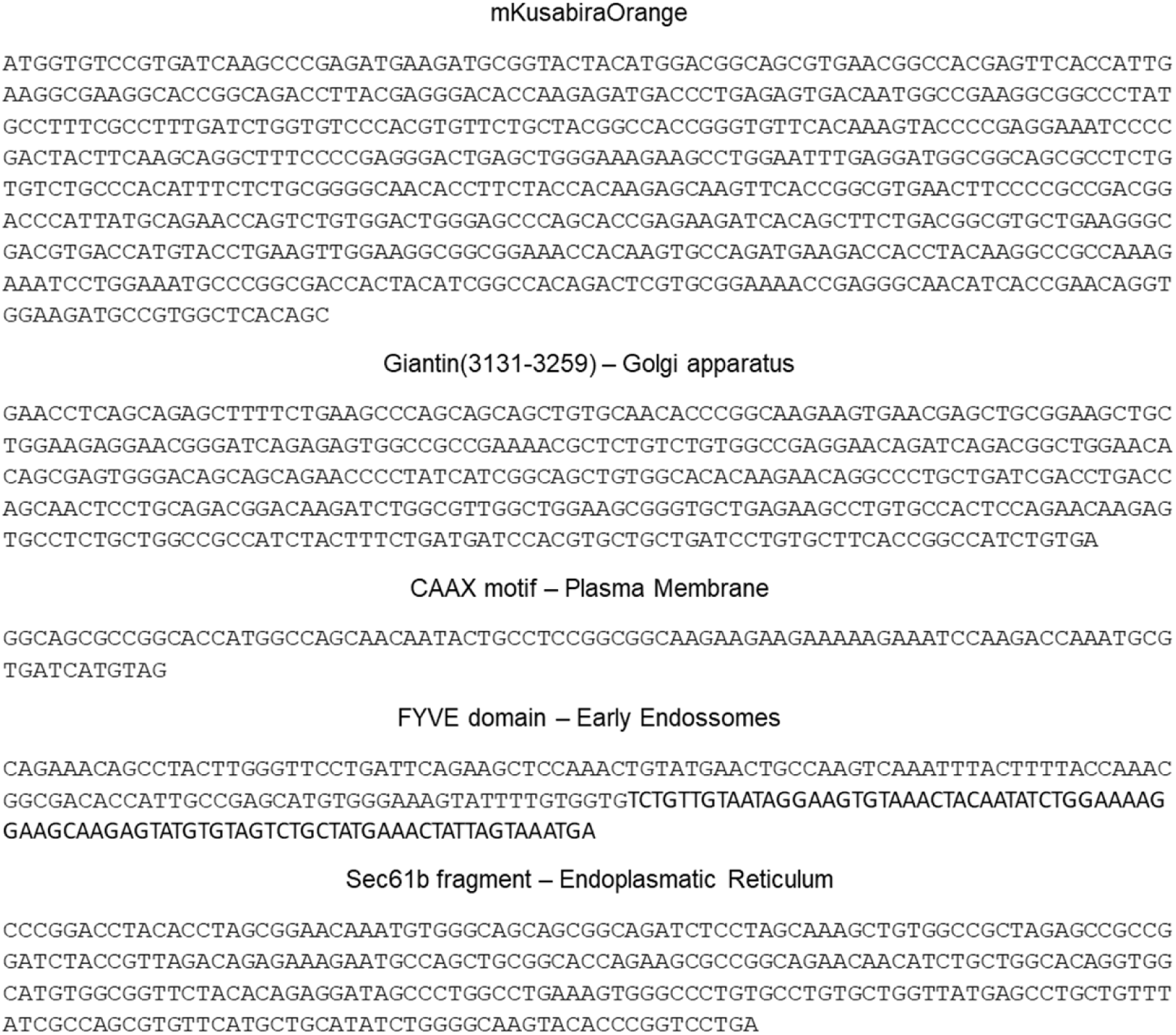
DNA coding sequences of plasmids used for microscopy experiments. Monomeric Kusabira-Orange2 (mKO) CDS was subcloned into a mammalian expression vector, pSF-CMV-AMP. Then, each CDS for K-Ras-derived CAAX motif (CAAX), FYVE domain of endofin (FYVE), Golgi targeting domain of giantin (Giantin) and the C-terminus of SecG (Sec61b) were fused to the 5’ end of mKO2 using Gibson assembly.

